# A heuristic model of the effects of phenotypic robustness in adaptive evolution

**DOI:** 10.1101/2020.04.19.048793

**Authors:** Emanuele Rigato, Giuseppe Fusco

## Abstract

A recent theoretical, deterministic model of the effects of phenotypic robustness on adaptive evolutionary dynamics showed that a certain level of phenotypic robustness (critical robustness) is a necessary condition for adaptation to occur and to be maintained in the course of evolution in most real organismal systems. We built an individual-based heuristic model to verify the soundness of these theoretical results through computer simulation, testing expectations under a range of scenarios for the relevant parameters of the evolutionary dynamics. These include the mutation probability, the presence of stochastic effects, the introduction of environmental influences and the possibility for some features of the population (like selection coefficients and phenotypic robustness) to change themselves during adaptation. Overall, we found a good match between observed and expected results, even for evolutionary parameter values that violate some of the assumptions of the deterministic model, and that robustness can itself evolve. However, from more than one simulation it appears that very high robustness values, higher than the critical value, can limit or slow-down adaptation. This possible trade-off was not predicted by the deterministic model.

## 1. Introduction

Phenotypic robustness is the quality of a biological system to maintain its phenotype in spite of internal (e.g., genetic) or external (e.g., environmental) modifications (Wagner, 2011; Klingenberg, 2019). This quality is widespread in living systems and can be observed at different levels of biological organization, from molecules to whole organisms (Kitano, 2004; Stelling et al., 2004; Wagner, 2005). Since phenotypic robustness to genetic mutations reduces the rate of appearance of favourable phenotypic mutations, at the level of the organism this might seem a trait that hinder adaptation by natural selection, by reducing the available variation for directional natural selection to act on. However, contrarily to this expectation, deriving from the focus of classic population genetics theory on adaptation from new mutations (Barrett and Schluter, 2008), recent theoretical work and several experimental studies on macromolecules have provided evidence for the capacity of this feature of the organism’s genotype-phenotype (G-P) map to enhancing adaptation (Gibson and Reed, 2008; Wagner, 2008; Rodrigues and Wagner, 2009; Draghi et al., 2010; Hayden et al., 2011; Barve and Wagner, 2013; Stiffler et al., 2015; Wei and Zhang, 2017). Even more recently, experimental evolution studies on *E. coli* have shown that phenotypic robustness, through the accumulation of cryptic genetic variation, can facilitate adaptation at the level of a whole organism (Rigato and Fusco, 2016; Zheng et al., 2019). Phenotypic robustness can foster adaptation through the accumulation of cryptic genetic variation (Wagner, 2012), so that in a new environment, phenotypically unexpressed genetic variation reveal accidentally “pre-adapted”, or “exapted” variants (Hayden et al., 2011), or by means of the scattering of genotypes with the same phenotype through the genotype space, which allows the population to access a greater number of new phenotypes upon mutation, increasing the probability of finding phenotypes that happen to have higher fitness (Rodrigues and Wagner, 2009; Zheng et al., 2019). In addition, robustness can support the spread of already present favourable phenotypic variants by dampening the probability of mutation, that in practice increases the evolutionary stability of favourable phenotypic variants (Rigato and Fusco, 2019).

In a recent study (Rigato and Fusco, 2019), we elaborated on a classical mathematical formalizations of evolutionary dynamics, the quasispecies model (Eigen at al., 1989), and showed that a certain level of phenotypic robustness is not only a favourable condition for adaptation to occur, but that it is also a necessary condition for short-term adaptation in most real organismal systems. This appears as a threshold effect, i.e. as a minimum level of phenotypic robustness (or, *critical robustness*) below which evolutionary adaptation cannot consistently occur or be maintained, even in the case of sizably selection coefficients and in the absence of any drift effect. These theoretical results, focusing on the instantaneous ability of the population to adapt, were obtained under the assumptions of some fixed evolutionary parameters (e.g., mutation rate, selection coefficient) and large, virtually infinite, population size. In other words, the analytical model was completely deterministic and concentrated on short-term effects.

We built and analysed an individual-based heuristic model to put to test the predictions of the deterministic model by Rigato and Fusco (2019) through computer simulation, verifying expectations under a range of scenarios for the relevant parameters of the evolutionary dynamics, some of which violate specific assumptions of the deterministic model. Parameters include the mutation probability, the presence of stochastic effects, the introduction of environmental influences and the possibility for some features of the population (such as selection coefficients and phenotypic robustness) to change themselves during adaptation.

## 2. The Model

Model description follows the ODD protocol for illustrating individual- and agent-based models (Grimm et al. 2006, 2010). The model was implemented in NETLOGO v. 5.0.3, (Wilensky, 1999; Tisue and Wilensky, 2004) and the scripts are available in the Supplementary material.

### 2.1. Purpose

The main purpose of the model is to explore the effects of phenotypic robustness in adaptive evolution, testing the influences of different sources of stochasticity (finite population size, environmental variation) with respect to the predictions of a deterministic model (Rigato and Fusco, 2019), which is a phenotypic version of the classic population genetics quasispecies model (Eigen et al., 1989). This is implemented by simulating phenotypic evolution of a finite population of entities under variable conditions.

### 2.2. Entities, state variables and timing

Entities of the model are asexual individuals. Each individual *i* has a genotype (not modelled) and a phenotype that includes its absolute fitness (number of offspring, *W*_*i*_) and phenotypic robustness (*ρ*_*i*_). Following Rigato and Fusco (2019), we adopted a narrow, quantitative definition of phenotypic robustness, that is the probability that, across one generation, mutation of a given genotype *g* takes to a genotype *g’* that exhibits the same phenotype of *g*, or, with other words, the probability that a mutation has no phenotypic effect. Different genotypes can map on the same phenotype and there are no fitness costs directly associated with phenotypic robustness. Depending on the simulation, state variables are: population mean relative fitness 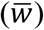 and mean robustness 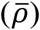. Relative fitness is computed with respect to a hypothetical maximum fitness value (*W*_*max*_), corresponding to a phenotypic optimum attainable in a given environment, thus *w*_*i*_=*W*_*i*_/*W*_*max*_. The model is time-discrete, one time step corresponding to one generation, and generations do not overlap.

### 2.3. Process overview

The process is schematically diagrammed in Figure 1. Each cycle (generation) starts with a parent population of *N* individuals. Parents reproduce according to their fitness value and die. Each single offspring can exhibit a mutated genotype with probability *η*, and, conditional on the mutated genotype, a mutated phenotype with probability 1-*ρ*. The magnitude of the phenotypic mutation can be fixed or modelled with a random variable, depending on the simulation. If the resulting offspring population is larger than *N*, this is reduced to size *N* through random elimination of the exceeding entities. Offspring of one generation are the parents of the succeeding generation. In simulation with changing environmental conditions, individual fitness values are modified before they reproduce.

**Figure 1.**
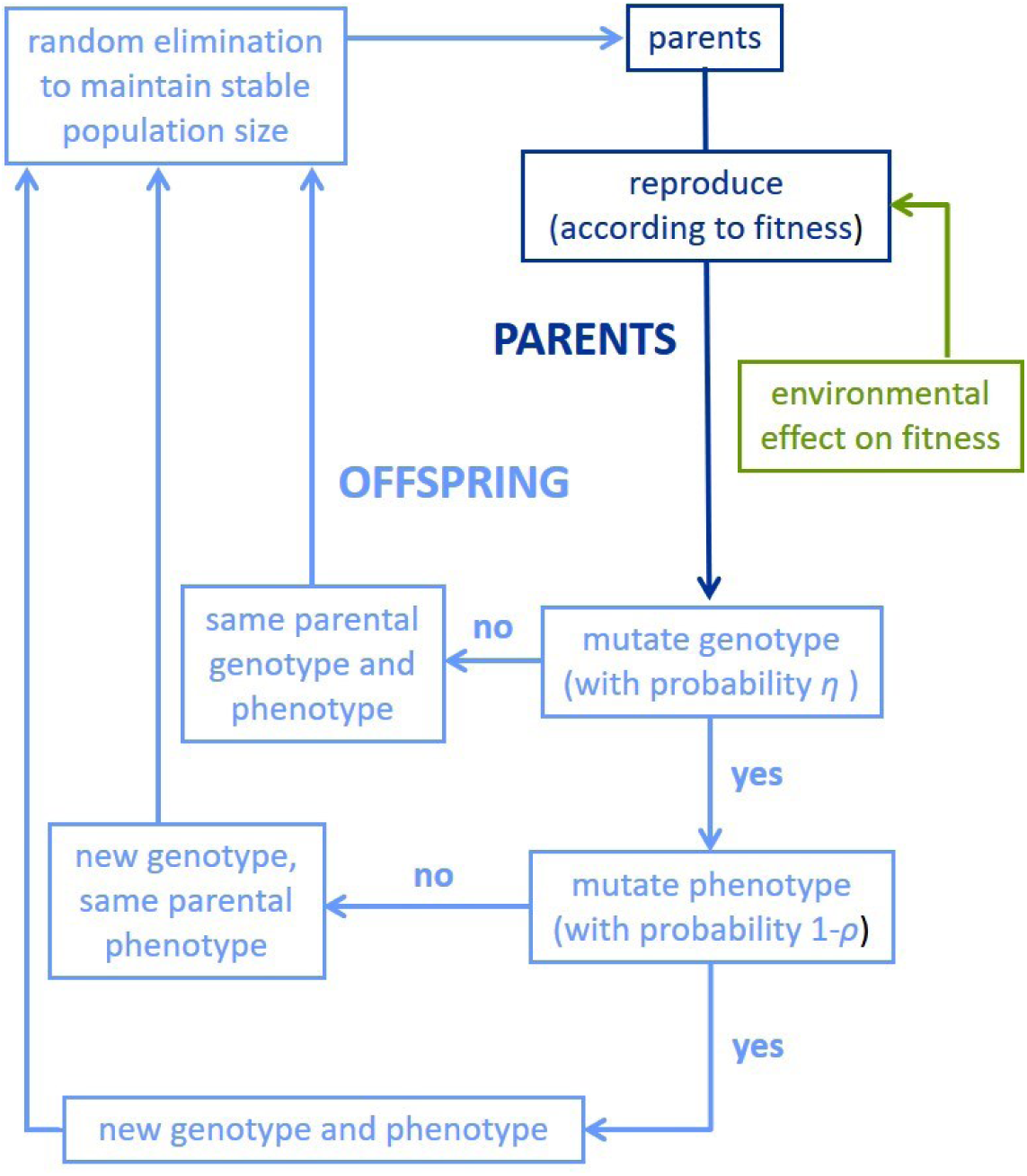
Schematic of the implemented individual-based model to study the effects of phenotypic robustness on the adaptive evolution in populations with finite size and under different evolutionary parameters. Individual organisms reproduce clonally and can mutate. The genotype-phenotype map, where multiple genotypes can map on the same phenotype, allows for genetic mutations with no phenotypic effects. Modifications in the environmental conditions (and thus in the shape of the adaptive landscape) are also considered.

### 2.4. Design concepts

Phenotypic evolution, measured as changes in the distribution of phenotypic values, is traced across generations in simulated populations. Population dynamics, in particular with respect to adaptation, emerge from the combined effects of heredity, genetic and phenotypic mutation (the latter, conditional on phenotypic robustness), natural selection (differential fecundity fitness among individuals) and demographic processes related to population size. Depending on the simulation, stochasticity may have effects on the occurrence of genotype and phenotype mutation, the magnitude of phenotype mutation, survival and environmental change.

### 2.5. Initialization

Parameters and their initial values depend on the simulation (see below).

### 2.6. Input

The model has no external input; parameters are updated according to internal rules of the model.

### 2.7. Sub-model “reproduction”

At each generation, each individual *i* clonally produces *W*_*i*_ offspring and dies.

### 2.8. Sub-model “mutation”

The genotype can mutate with probability *η* and if a genetic mutation occurs, the phenotype can mutate with probability (1-*ρ*). Depending on the simulation, mutation can affect only fitness or both fitness and phenotypic robustness. In the latter case, the new parameters of the individual are modified independently. Individual phenotypic values are updated according to the following rules:

- in simulations with fixed selection coefficient *s* (Simulations 1 and 2), the new mutated fitness in individual *i* is set to 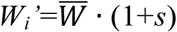. The sign of *s* can be positive with probability *τ*=0.17 and negative with probability 1-*τ*. Probability *τ* derives from assuming an optimal mutation’s phenotypic effect on the Fisher’s (1930) standardized scale (Orr, 2000). However, different values of *τ* only change the kinematics of the process, not its most salient feature in these simulations, i.e. the tendency towards either higher or lower values of mean fitness in proportion to *ρ*.
- in simulations with variable selection coefficients (Simulations 3 and 4), the new mutated fitness in individual *i* is set to a value ranging from 0 to the fitness maximum with equal probability, *W*_*i*_*’*=U[0, *W*_*max*_]. Accordingly, both the magnitude and probability of mutant fitness gain decrease as the population approaches a posited value of maximum fitness, as expected under Fisher’s (1930) *geometric model* (see also Orr, 1998, 2000), or, equivalently, on the basis of the more general *extreme value theory* (Orr, 2005).
- in the simulation with variable robustness (Simulation 4), the new mutated robustness value is set to a value ranging from 0 to 1 with equal probability, *ρ*_*i*_*’*=U[0, 1].

All these updating rules assume a very rugged fitness landscape, or equivalently, a totally uncorrelated G-P map, as upon the mutation the original fitness (or robustness) value of the individual does not influence the probability distribution of the new value. This is reasonable assumption in consideration of the high dimensionality of the genotype space (Gavrilets, 1997; Aguirre et al., 2018) and the non-linearity of the G-P map (Ahnert, 2017; Green et al., 2017; Sailer and Harms, 2017).

### 2.9. Sub-model “environmental change”

Environmental change is implemented as a negative effect on individual fitness (environmental deterioration, sensu Fisher, 1930). The magnitude of the negative effect is quantified as a parameter (*X*) that is subtracted to the individual absolute fitness values (*W*_*i*_) before reproduction. *X* is modelled as a discrete uniform random variable (U[a,b]), with a different range ([0, 0], [0, 2], [0, 4], or [0, 6]) and mean (0, 1, 2, or 3) in different runs. Thus, the effect of *X* combines environmental disturbance with environmental instability.

### 2.10. Sub-model “maximum population size”

At each generation, if offspring population is larger than *N*, the population is reduced to size *N* through random elimination of the exceeding entities. The surviving entities are going to be the parents of the succeeding generation.

## 3. Simulations

The deterministic model predicts a minimum level of phenotypic robustness for adaptation to occur, i.e. for the population mean fitness to increase. The value of this *critical robustness* (*ρ*_*c*_) depends on the magnitude of the selection coefficient (*s*) and on the genotype mutation probability (*η*) according to the following equation

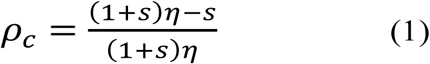

To test the soundness of this model under different combinations of stochastic effects, we run four simulations. Simulations went on for 100-500 generations. Each simulation included several runs, each characterized by different initialization parameters, and a number of replicas for each run. Parameters were explored in ranges considered particularly challenging for the model, i.e. for values that significantly departed from the assumptions of the deterministic model, such as quite small population sizes, high selection coefficients and variable selection regimes. Values of mutation probability, instead, were chosen in line with observed data in real organisms.

## 4. Results

### 4.1. Existence of *ρ*_*c*_ for populations of finite size: Simulation 1

This simulation aimed at testing the existence of a minimum level of phenotypic robustness below which the population mean fitness (the state variable) does not increase significantly across time (generations). This was implemented by fixing the selection coefficient *s* while exploring various parameters combinations of population size *N*, genotype mutation probability *η* and phenotypic robustness *ρ*. Adaptation was said to have occurred when a significant increase in the population mean relative fitness (one-tail Student’s t-test, α=0.05) was recorded within the 500 generations.

- Fixed parameters (equal among runs): *s*=0.1; *W*_*max*_=30
- Fixed parameters (different among runs): *η*=0.8, 0.9, 1.0; *N*=50, 100, 500, 1000; *ρ*=0.70, 0.75, 0.80, 0.85, 0.90, 0.95, 0.99
- Internal variables: *W*_*i*_ (initial value=10 for all individuals), 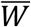
- State variable: 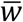
- Generations: 500
- Runs: 84 (3×4×7; *η, N, ρ* values, respectively)
- Replicas: 10 per run

#### Observations

The 84 runs (Fig. 2) show no adaptation for *N*=50, and for *N*=100 there is adaptation only at *ρ*=0.99. For larger-size populations (*N*=500 and 1000), adaptation occurs consistently for *ρ*≥0.90. Thus, the robustness threshold at these population sizes seems to be located in between 0.85 and 0.90, a range of values very close to that expected by the deterministic model (0.89-0.91). As a general pattern, adaptation occurs more consistently to the increase of both *η* and *N*, although the latter seems to be a more influential parameter. In parallel, the higher the robustness, the faster the adaptation, when it occurs, or the slower and more limited the decrease in the mean fitness when adaptation does not occur. Among the larger-size populations (*N*=500 and 1000), those with the highest robustness *ρ*=0.99 are often outperformed by populations with *ρ=*0.95 and 0.90. It seems that while robustness above *ρ*_*c*_ enhances adaptation in proportion to its value, there is an upper limit of *ρ* above which the boosting effect declines. This effect was not anticipated on the basis of the deterministic model.

**Figure 2.**
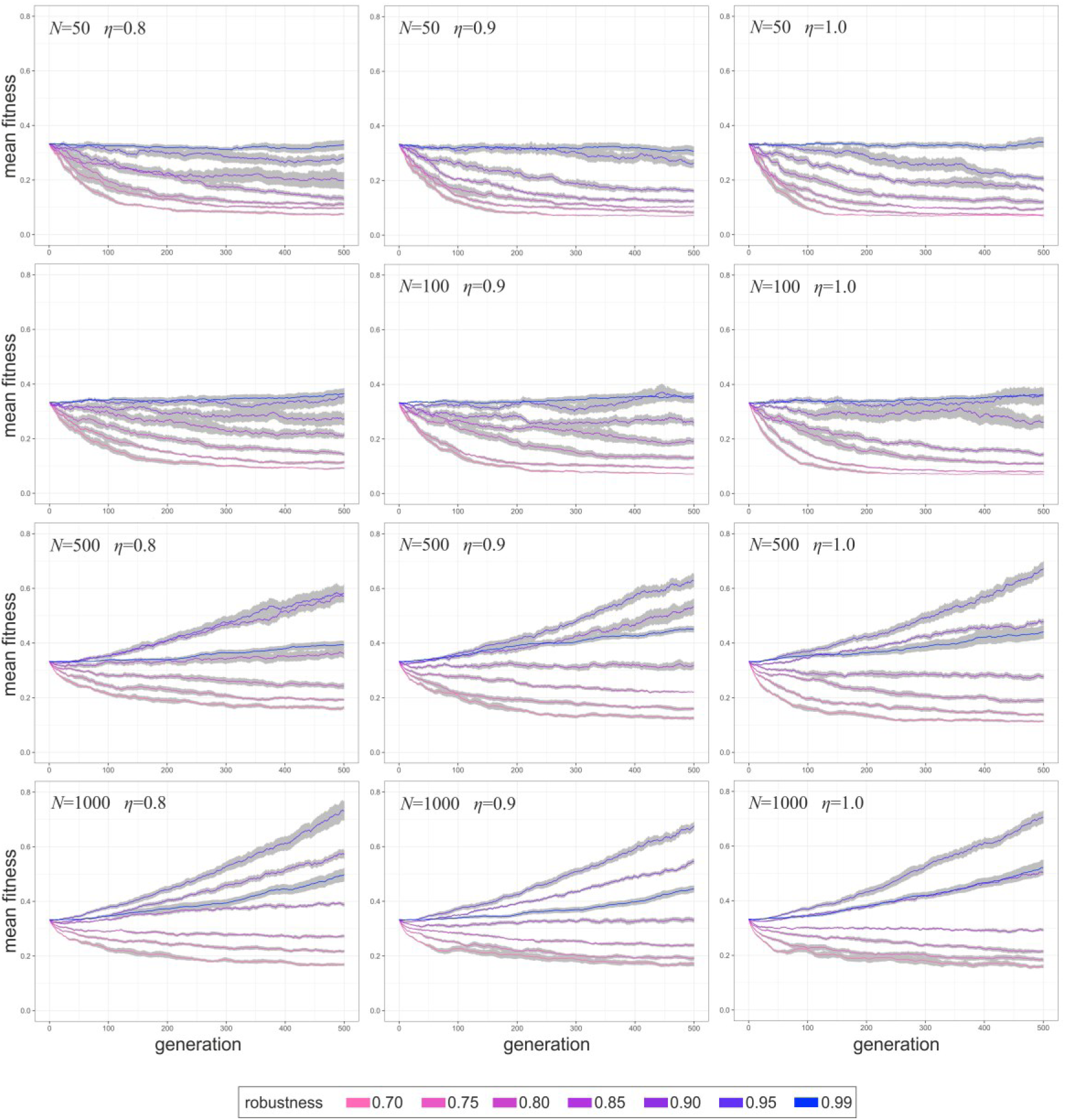
Adaptation dynamics in populations with different robustness (*ρ*), under fixed selection coefficient (*s*=0.1) and with different population size (*N*) and genotype mutation probability (*η*) during 500 generations (Simulation 1). Lines are means over 10 replicas and the grey areas show the standard error of the means. Adaptation is stated to have occurred if at the end of the 500 generations the populations reached a mean relative fitness significantly higher (p<0.05) than the initial fitness value, set at 1/3 of the of the relative fitness with respect to an optimal phenotype (*w*=1).

Finally, as expected by the deterministic model, at *N*=1000 the critical robustness for a mutation probability of *η*=0.8 is lower than that at *η*=0.9 and *η*=1.0, so that at *η*=0.8 we observe adaptation even at *ρ*=0.85. This effect is visible also at *N*=500, although adaptation at *η*=0.8 and *ρ*=0.85 is statistically not significant (*p*=0.08) after 500 generations.

Overall, what can be observed from Simulation 1 is that phenotypic robustness enhances adaptation rate in populations with *ρ>ρ*_*c*_ and buffers the population mean fitness decrease in populations with *ρ<ρ*_*c*_.

### 4.2. Value of *ρ*_*c*_ in populations of finite size: Simulation 2

Simulation 2 adopted the same design of Simulation 1, but recorded the minimum level of robustness below which adaptation was not maintained or did not increase (one-tail Student’s t-test, α=0.05, on fitness at generation 500) in runs with variable *ρ*, under different combinations of *s* and *η* in populations with nearly constant finite size *N*. The minimal value for the parameter *s* was set to 0.02, so to have *sN*≥1, however selection must be considered week with respect to random drift for the smallest values in the parameter interval. Observed *ρ*_*c*_ were compared with the values predicted by the deterministic model.

- Fixed parameters (equal among runs): *N*=500; *W*_*max*_=30
- Fixed parameters (different among runs): *η*=0.001, 0.10, 0.20, 0.30, 0.40, 0.50, 0.60, 0.70, 0.80, 0.90, 0.99; *s*=0.02, 0.03, 0.04, 0.05, 0.06, 0.07, 0.08, 0.09, 0.10; *ρ*=0.001, 0.10, 0.20, 0.30, 0.40, 0.50, 0.60, 0.70, 0.80, 0.85, 0.90, 0.95, 0.99
- Internal variables: *W*_*i*_ (initial value=10 for all individuals), 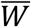
- State variable: 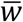
- Generations: 500
- Runs: 1287 (11×9×13; *η, s, ρ* values, respectively)
- Replicas: 10 per run

#### Observations

Overall, the 99 *ρ*_*c*_ values obtained through simulation are fairly close to the corresponding values on the surface of the expected *ρ*_*c*_ in the parameter space defined by *s* and *η* under equation (1) (Fig. 3). The deterministic model explains a large fraction of the observed variance in *ρ*_*c*_ (r^2^=95.08%, n=99). In general, there is a tendency for the predicted values to be slightly higher than the observed, with residuals declining with increasing *η*. This depends on the fact than in the simulation, at variance with the deterministic model, new mutations are continuously introduced, with a transient lowering effect on the average fitness at every generation before selection. This slightly amplifies the *s* values, allowing adaptation at smaller *ρ* than expected, in particular at small *η* (see equation (1)).

**Figure 3.**
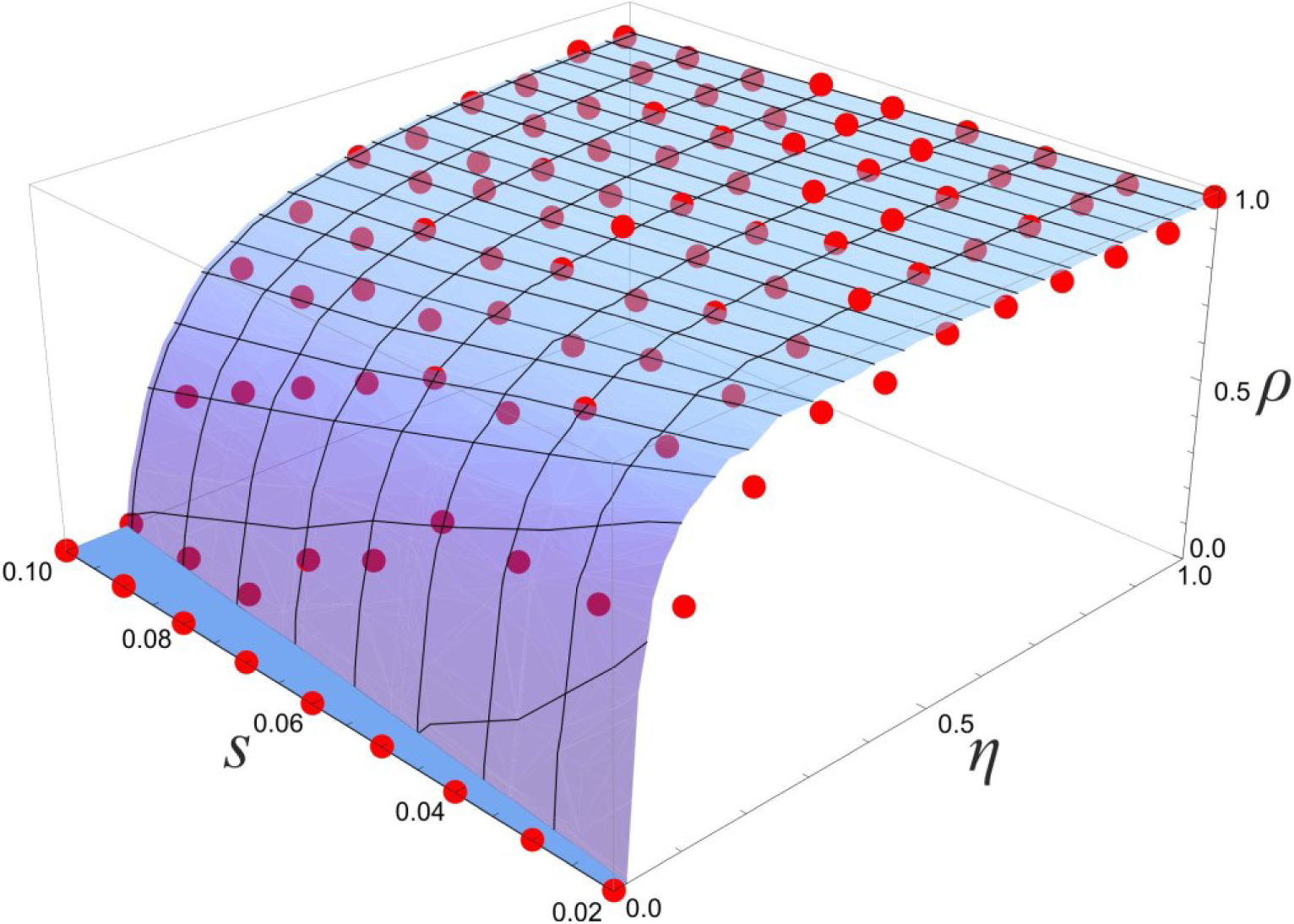
Values of critical robustness (*ρ*_*c*_) in populations of size *N*=500, with different mutation probability (*η*) and different fixed selection coefficients (*s*) (Simulation 2). The observed critical robustness (red dots) is the minimum value of robustness (among the set of values 0.001, 0.10, 0.20, 0.30, 0.40, 0.50, 0.60, 0.70, 0.80, 0.85, 0.90, 0.95, 0.99) for which adaptation has occurred or has been maintained after 500 generations, i.e. for which the populations scored a final mean fitness not significantly below the initial fitness value (p<0.05). Critical robustness values obtained through simulation are to be compared with the corresponding points on the surface of the expected *ρ*_*c*_ values derived by the deterministic model.

### 4.3. Effect of *ρ* on the limits to adaptation: Simulation 3

Simulation 3 aimed at testing the maximum population mean fitness that is reachable in finite size populations with different levels of robustness. Here, both the magnitude and the probability of fitness gain in the mutants decrease as the population approaches the maximum average fitness, as entailed by Fisher’s (1930) geometric model (Waxman and Welch, 2005).

- Fixed parameter (equal among runs): *W*_*max*_=30
- Fixed parameters (different among runs): *η*=0.8, 0.9, 1.0; *N*=50, 100, 500, 1000; *ρ*=0.70, 0.75, 0.80, 0.85, 0.90, 0.95, 0.99
- Internal variables: *W*_*i*_ (initial value=15 for all individuals), 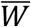
- State variable: 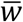
- Generations: 500
- Runs: 84 (3×4×7; *η, N, ρ* values, respectively)
- Replicas: 10 per run

#### Observations

In all runs (Fig. 4), the level of adaptation reaches a plateau, whose height with respect to the maximum value attainable is proportional to the level of robustness, *ρ*. The amplitude of the oscillations around the plateau value decreases with both the population size, *N*, and with the robustness, *ρ*. At variance with Simulation 1, there is no declining fitness in populations with low robustness because population can settle on a plateau (at a given distance from the fitness maximum) which grants sufficiently high coefficients of selection for any *ρ* value. The initially very high selection coefficients explain the fast achievement of the plateau in all runs. The probability of mutation, *η*, does not visibly affect the elevation of the plateaus, whereas the population size, *N*, has the effect of slightly lifting the plateaus. Although the relative height of the plateaus reached in each run is significant for our argument, with the higher elevations attained by population with higher *ρ*, their absolute values are not, because they depend on the specific modelling of the variation of the coefficients of selection as adaptation progress (in our heuristic model, a sampling from a uniform distribution, see Section 2.8).

**Figure 4.**
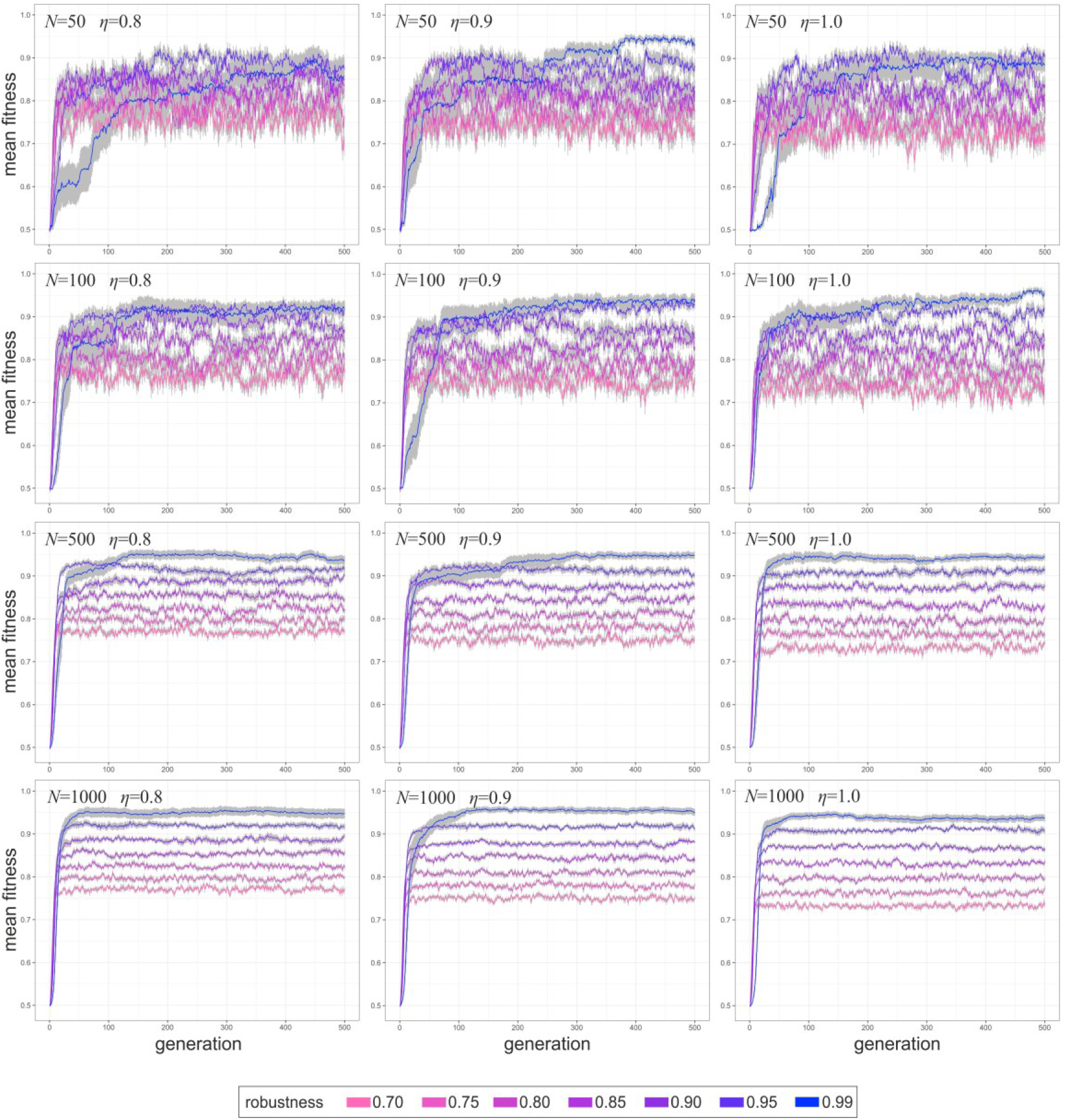
Adaptation dynamics in populations with different robustness (*ρ*), under declining selection coefficient (*s*) according to Fisher’s geometric model and with different population size (*N*) and genotype mutation probability (*η*) during 500 generations (Simulation 3). Lines are means over 10 replicas and the grey areas show the standard error of the means. Initial average fitness values is set at 1/2 of the of the relative fitness with respect to an optimal phenotype (*w*=1).

Like in Simulation 1, adaptive dynamics with the highest robustness, *ρ*=0.99, show a behaviour slightly uneven with respect to lower robustness values. Although simulations with *ρ*=0.99 reach the highest plateaus, the rise to the plateau is slightly retarded with respect to simulations with lower *ρ*. This effect was not predicted by the deterministic model.

### 4.4. Robustness evolvability: Simulation 4

This simulation aimed at probing the evolvability of phenotypic robustness itself, under different environmental conditions. This was implemented by fixing both genotype mutation probability *η* and population size *N*, while exploring the effect of different levels of environmental deterioration and instability, represented by the random variable *X*. As in Simulation 3, both the magnitude and the probability of fitness gain in the mutants decrease as the population approaches the maximum average fitness. However, at variance with all the preceding simulations, phenotypic robustness itself could vary among individuals and evolve across generations. Population mean phenotypic robustness thus adds to population mean relative fitness as a new state variable.

- Fixed parameters (equal among runs): *η*=1.0; *W*_*max*_=30; *N*=1000
- Fixed parameters (different among runs): *X*=0, U[0, 2], U[0, 4], U[0, 6] (with 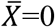, 1, 2, 3, respectively)
- Internal variables: *W*_*i*_ (initial value=10 for all individuals); *ρ*_*i*_ (initial value=0.1 for all individuals)
- State variables: 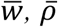
- Generations: 100
- Runs: 4
- Replicas: 10 per run

#### Observations

The higher the level of environmental deterioration 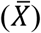, the lower the mean fitness value that a population can reach (Fig. 5). A similar pattern is observable in the evolution of robustness, which increases slower and reaches lower values in more degraded environments.

**Figure 5.**
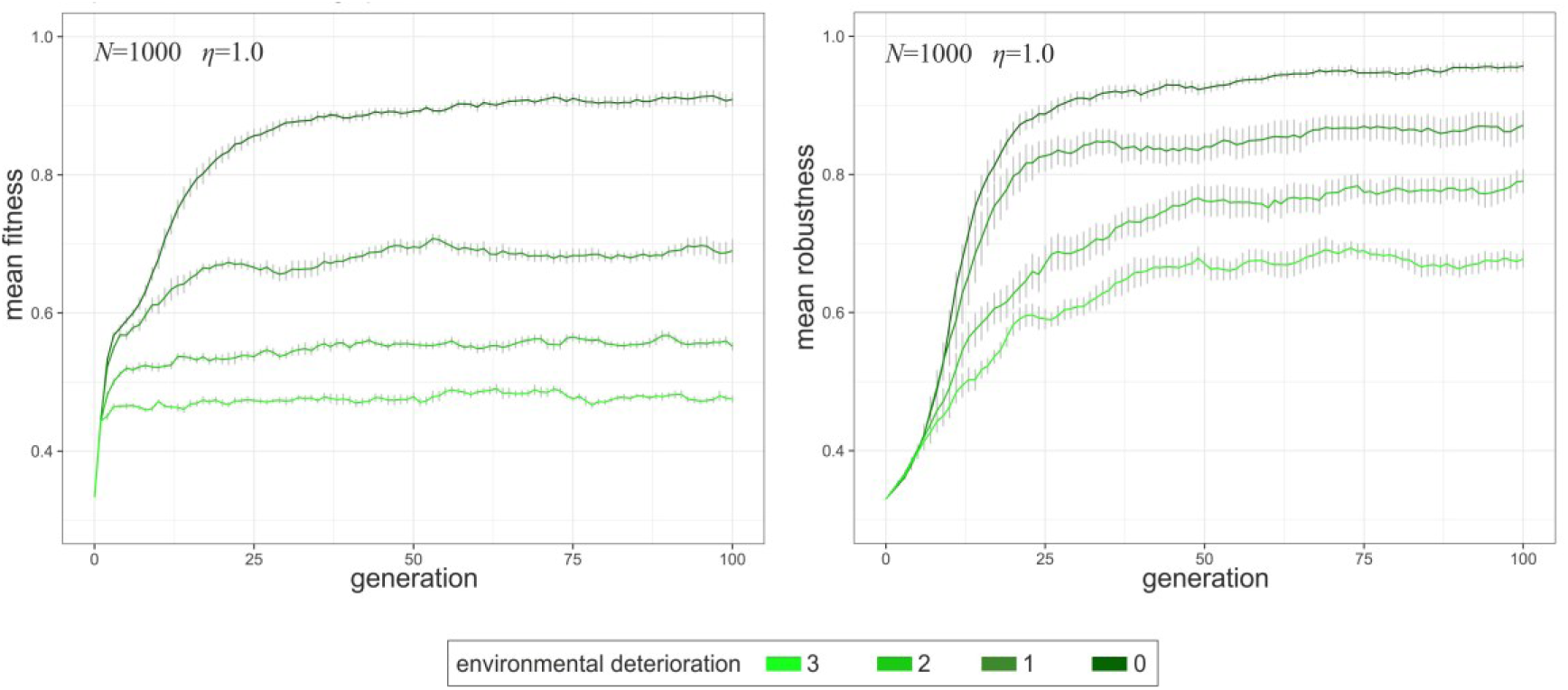
Evolution of fitness (*w*, left) and phenotypic robustness (*ρ*, right) in populations of size *N*=1000 and with mutation probably *η*=1, under different level of average environmental deterioration during 100 generations (Simulation 4). Lines are means over 10 replicas and bars are the standard error of the means.

During adaptation, natural selection promotes a simultaneous increase in fitness and robustness toward a maximum value attainable for the specific level of environmental deterioration. However, while for 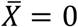, population mean fitness and mean robustness increase in parallel until generation 100, their values at each generation closely matching the combinations 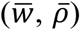 derived through Simulation 3 (with *N*=1000 and *η*=1.0; Fig. 4), in case of environmental deterioration 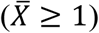 fitness reaches a limit value faster than robustness.

The main result of this simulation is to display the evolvability of robustness itself, under a range of different environmental conditions. However, the simulation also exposes the limits that the environment can pose to adaptation. Very clearly, and expectedly, these appear as constraints to the maximum fitness attainable 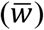. But beyond that, it is also apparent that very high robustness is less favourable in degraded and unstable environments and that under the same conditions robustness has a less marked positive effect on fitness.

## 5. Discussion

Through computer simulation, we studied the effects of phenotypic robustness in the evolution of populations of different size and under different evolutionary parameters, and compared the results of the simulations with the predictions of a deterministic model by Rigato and Fusco (2019).

In agreement with the theoretical model, we show that a minimum level of phenotypic robustness is required for adaptation to occur, also in populations subject to the stochastic effects of a relatively small finite size (Simulation 1), and that these observed critical values of robustness are close to those derived by the deterministic model for a large part of the parameter space (Simulation 2). Although in populations of very small size (*N*≤100), in practice there is no *ρ* value able to effectively contrast the dominant effect of drift (Ohta, 1992), even for the very large selection coefficient modelled (*s*=0.1), nonetheless, different levels of robustness may have an effect on evolutionary dynamics irrespective of adaptation: phenotypic robustness higher than *ρ*_*c*_ enhances adaptation rate, whereas *ρ* below those values buffers the progressive loss in population mean fitness in proportion to *ρ*. The implications of considering the quasispecies model in the context of finite-size populations have been explored, although with different focuses, in other analytical and numerical analyses (Park et al., 2010; Lorenz et al., 2013). Among other results, these works showed that time-averaged fitness is expected to be lower for finite populations than it is for infinite populations. Our results are silent on these long-term effects, since Simulations 1 and 2 were designed to explore the kinematics of adaptation under different robustness conditions, rather than its ‘final’ achievement. However, they are nonetheless compatible with these previous results.

The enhancing effect of robustness on adaptation appears clearly also when considering a reduction in the average fitness gain upon mutation as the population approaches the maximum average fitness (Fisher’s geometric model, Simulation 3) and when introducing negative effects on fitness due to environmental deterioration and instability (Simulation 4). In both cases, the level of adaptation reaches value directly proportional to the level of robustness. There are however limits to the maximum attainable fitness with respect to a theoretical optimum. This is because the phenotypic robustness needed for a given advantageous phenotype to spread throughout the population is inversely related to its selective advantage (*s*) in any given moment, as equation (1) shows. While the population mean fitness get closer to the optimum, the selection coefficients become progressively smaller, until a point where the population can no longer increase its fitness, because *ρ* falls below the critical value needed for adaptation.

Alongside these computational confirmations of the deterministic model, there are effects of robustness on adaptation that were not predicted by it. Simulations 1, 3 and 4 show that very high robustness values, beyond the critical value, can limit or slow-down adaptation. These divergences can stem from several features that distinguish the theoretical model from the computational model, but one appears to be especially relevant. Among the different ways in which phenotypic robustness can influence adaptation, Rigato and Fusco (2019) deterministic model concentrated on the competence of robustness to support the spread of already present favourable phenotypic variants, while evidence of the enhancing effect of robustness on adaptation produced by other means, e.g. through the accumulation of cryptic genetic variation, was provided by other studies (Hayden et al., 2011; Rigato and Fusco, 2016; Zheng et al., 2019). However, despite our simulations do not explicitly contemplate the modelling of the evolution of the genetic structure of the population, some aspects of longer-term evolutionary dynamics were inevitably introduced. Among these, and most relevant here, is the probability that new favourable phenotypic mutations appear in the population during the process of adaptation. In simulated populations, this introduces a trade-off between the positive effect of robustness in facilitating the spread of favourable mutations, and the negative effect of making the appearance of such mutations less frequent. This trade-off was not contemplated in the deterministic model, but it can be of relevance in real evolutionary dynamics. Although standing variation may be of paramount importance in the process of adaptation (Matuszewski et al., 2015; Lai et al., 2019), undoubtedly there are circumstances where adaptation critically depends on a supply of de-novo mutations (Exposito-Alonso et al., 2018).

The effects of environmental changes during adaptation were completely outside the scope of deterministic model by Rigato and Fusco (2019). However, the results of the simulations can be compared with the findings of other studies on the subject, both analytical and computational. With regard to fitness, our observation that high levels of environmental instability limit the population mean fitness is in agreement with the predictions of many genetic models, which show that frequent environmental changes prevent populations from reaching a fitness peak (e.g., Fisher, 1930; Trubenová et al., 2019). With regards to robustness, several theoretical works have addressed the question of under what conditions natural selection would lead to an increased robustness, but the results they obtained are not all congruent or easily comparable. For instance, Felix and Wagner (2008) argued that from theoretical literature (see references therein) it emerges that high robustness tends to evolve more easily when perturbations (either genetic or environmental) are abundant. We observed robustness to increase in all the conditions we tested, and with respect to the level of environmental disturbance we found the opposite relationship. While in a stable environment, and possibly in an environment that changes slowly and monotonically through time, phenotypic robustness evolves to reach values close to 1, in case of a repeatedly and unpredictably changing environment robustness does not reach such high values. However, it should be noted that in our simulation the evolutionary change in robustness is accomplished by modifying a single feature of the genotype-phenotype map, that is the fraction of genotypes that map on the same phenotype. This does not exclude that other qualities of robustness, like the capacity to accumulate cryptic genetic variation, may flourish under different environmental conditions. A different prediction was formulated by Hordijk and Altenberg (2020), by analysing a computational model for the evolution of ontogeny based on cellular automata. They showed that developmental systems evolving high modularity tend also to evolve mutational robustness, while systems where the production of a phenotype depends on complex interaction of many contributing components do not. We did not explicitly modelled development in our simulations, but, like in Rigato and Fusco (2019), we assumed a G-P map with extended pleiotropy (*ubiquitous pleiotropy*; Visscher and Yang, 2016) and with almost every trait affected by many genes (*omnigenic model*; Boyle et al., 2017). Thus we can probably state that our simulations showed the occurrence of robustness evolution far from the case of modularity.

Our results are more in agreement with the arguments of de Visser et al. (2003), who claimed that high mutation rates, large populations, and asexual reproduction (three conditions matched by our model) generally favour the evolution of *adaptive robustness*, i.e. a form of robustness where the buffering of phenotype with respect to some source of variation has been a direct target of natural selection, rather than having been an indirect by-product of selection on other traits (*intrinsic robustness*). While we did not probe the evolution of robustness in the opposite conditions, our simulation clearly produce the evolution of adaptive robustness, because fitness cannot increase if not accompanied by an increase of robustness.

In conclusion, these simulations, which prove the positive effects of robustness on adaptation under less idealistic conditions with respect to those assumed by the theoretical model, and show the evolvability of robustness along with different levels of environmental perturbation, can contribute to explain why robustness is so common in living systems.

## Supporting information

Simulation scripts

## Acknowledgements

This work has been supported by a grant from the Italian Ministry of Education, University and Research (MIUR) to GF.

## Notes

### Competing Interest Statement

The authors have declared no competing interest.

## References

Aguirre J., Catalán, P., Cuesta, J. A. and Manrubia, S. (2018). On the networked architecture of genotype spaces and its critical effects on molecular evolution. Open Biology, 8: 180069.

Ahnert, S. E. (2017). Structural properties of genotype–phenotype maps. Journal of The Royal Society Interface, 14: 20170275.

Barrett R. D and Schluter D. (2008). Adaptation from standing genetic variation. Trends in Ecology and Evolution, 23: 38–44.

Barve, A. and Wagner, A. (2013). A latent capacity for evolutionary innovation through exaptation in metabolic systems. Nature, 500: 203–206.

Boyle, E. A., Li, Y. I., and Pritchard, J. K. (2017). An expanded view of complex traits: from polygenic to omnigenic. Cell, 169: 1177–1186.

de Visser, J. A. G. M., Hermisson, J., Wagner, G. P., Ancel Meyers, L., Bagheri-Chaichian, H., Blanchard, J. L., Chao, L., Cheverud, J. M., Elena, S. F., Fontana, W., Gibson, G., Hansen, T.F., Krakauer, D., Lewontin, R.C., Ofria, C., Rice, S. H., von Dassow, G., Wagner, A. and Whitlock, M. C. (2003). Perspective: Evolution and detection of genetic robustness. Evolution, 57: 1959–1972.

Draghi, J. A., Parsons, T. L., Wagner, G. P. and Plotkin, J. B. (2010). Mutational robustness can facilitate adaptation. Nature, 463: 353–355.

Eigen, M., McCaskill, J., and Schuster, P. (1989). The molecular quasi-species. Advances in Chemical Physics, 75: 149–263.

Exposito-Alonso, M., Becker, C., Schuenemann, V. J., Reiter, E., Setzer, C., Slovak, R, Brachi, B., Hagmann, J., Grimm, D. G., Chen, J., Busch, W., Bergelson, J., Ness, R. W., Krause, J., Burbano, H. A. and Weigel, D. (2018). The rate and potential relevance of new mutations in a colonizing plant lineage. PLoS Genetics, 14: e1007155.

Félix, M.-A. and Wagner, A. (2008). Robustness and evolution: concepts, insights and challenges from a developmental model system. Heredity, 100: 132–140.

Fisher, R. A. (1930). The Genetical Theory of Natural Selection. Oxford University Press, Oxford.

Gavrilets, S. (1997). Evolution and speciation on holey adaptive landscapes. Trends in Ecology and Evolution, 12: 307–312.

Green, R. M., Fish, J. L., Young, N. M., Smith, F. J., Roberts, B., Dolan, K., Choi, I., Leach, C. L., Gordon, P., Cheverud, J. M., Roseman, C. C., Williams, T. J., Marcucio, R. S. and Hallgrímsson, B. (2017). Developmental nonlinearity drives phenotypic robustness. Nature Communications, 8: 1970.

Grimm, V., Berger, U., Bastiansen, F., Eliassen, S., Ginot, V., Giske, J., Goss-Custard, J., Grand, T., Heinz, S. K., Huse, G., Huth, A., Jepsen, J. U., Jørgensen, C., Mooij, W. M., Müller, B., Pe’er, G., Piou, C., Railsback, S. F., Robbins, A. M., Robbins, M. M., Rossmanith, E., Rüger, N., Strand, E., Souissi, S., Stillman, R. A., Vabø, R., Visser, U. and DeAngelis, D.L. (2006). A standard protocol for describing individual-based and agent-based models. Ecological Modelling, 198: 115–126.

Grimm, V., Berger, U., DeAngelis, D. L., Polhill, J. G., Giske, J. and Railsback, S. F. (2010). The ODD protocol: a review and first update. Ecological Modelling, 221: 2760–2768.

Gibson, G. and Reed, L. K. (2008). Cryptic genetic variation. Current Biology, 18: R989–R990.

Hayden, E. J., Ferrada, E. and Wagner, A. (2011). Cryptic genetic variation promotes rapid evolutionary adaptation in an rna enzyme. Nature, 474: 92–95.

Hordijk, W. and Altenberg, L. (2020). Developmental structuring of phenotypic variation: A case study with a cellular automata model of ontogeny. Evolution and Development, 22: 20–34.

Kitano, H. (2004). Biological robustness. Nature Reviews Genetics, 5: 826–837.

Klingenberg, C.P. (2019). Phenotypic plasticity, developmental instability, and robustness: The concepts and how they are connected. Frontiers in Ecology and Evolution, 7: 56.

Lai, Y.-T., Yeung, C. K. L., Omland, K. E., Pang, E.-L., Hao, Y. U., Liao, B.-Y., Cao, H.-F., Zhang, B.-W., Yeh, C.-F., Hung, C.-M., Hung, H.-Y., Yang, M.-Y., Liang, W., Hsu, Y.-C., Yao, C.-T., Dong, L., Lin, K. and Li, S.-H. (2019). Standing genetic variation as the predominant source for adaptation of a songbird. Proceedings of the National Academy of Sciences of the United States of America, 116: 2152–2157.

Lorenz, D. M., Park, J.-M., and Deem, M.W. (2013). Evolutionary processes in finite populations. Physical Review, E 87: 022704.

Matuszewski, S., Hermisson, J., and Kopp, M. (2015). Catch me if you can: adaptation from standing genetic variation to a moving phenotypic optimum. Genetics, 200: 1255–1274.

Orr, H. A. (1998). The population genetics of adaptation: the distribution of factors fixed during adaptive evolution. Evolution, 52: 935–949.

Orr, H. A. (2000). Adaptation and the cost of complexity. Evolution, 54: 13–20.

Orr, H. A. (2005). The genetic theory of adaptation: a brief history. Nature Review Genetics, 6: 119–127.

Ohta, T. (1992). The nearly neutral theory of molecular evolution. Annual Review of Ecology and Systematics, 23: 263–286.

Park J.-M., Muñoz E., and Deem M.W. (2010). Quasispecies theory for finite populations. Physical Review, E 81: 011902.

Rigato, E. and Fusco, G. (2016). Enhancing effect of phenotype mutational robustness on adaptation in *Escherichia coli*. Journal of Experimental Zoology Part B: Molecular and Developmental Evolution, 326: 31–37.

Rigato, E. and Fusco, G. (2019). Effects of phenotypic robustness on adaptive evolutionary dynamics. bioRxiv, 511691 (doi:10.1101/511691).

Rodrigues, J. F. M. and Wagner, A. (2009). Evolutionary plasticity and innovations in complex metabolic reaction networks. PLoS Computational Biology, 5: e1000613.

Sailer, Z.R. and Harms, M.J. (2017). Detecting high-order epistasis in nonlinear genotype-phenotype maps. Genetics, 205: 1079–1088

Stelling, J., Sauer, U., Szallasi, Z., Doyle, F. J., and Doyle, J. (2004). Robustness of cellular functions. Cell, 118: 675–685.

Stiffler, M.A., Hekstra, D.R. and Ranganathan, R. (2015). Evolvability as a function of purifying selection in TEM-1 beta-lactamase. Cell, 160: 882–892.

Tisue, S. and Wilensky, U. (2004). Netlogo: A simple environment for modeling complexity. In International conference on complex systems, vol. 21, pp. 16–21.

Boston, MA. Trubenová, B. Krejca, M. S., Lehre, P. K. and Kötzing, T. (2019). Surfing on the seascape: Adaptation in a changing environment. Evolution, 73: 1356–1374

Visscher, P. M. and Yang, J. (2016). A plethora of pleiotropy across complex traits. Nature Genetics, 48: 707.

Wagner, A. (2005). Distributed robustness versus redundancy as causes of mutational robustness. Bioessays, 27:176–188.

Wagner, A. (2008). Robustness and evolvability: a paradox resolved. Proceedings of the Royal Society of London B, 275: 91–100.

Wagner, A. (2011). The Origins of Evolutionary Innovations: A Theory of Transformative Change in Living Systems. Oxford University Press, Oxford.

Wagner, A. (2012). The role of robustness in phenotypic adaptation and innovation. Proceedings of the Royal Society of London B, 279: 1249–1258.

Waxman, D. and Welch, J. J. (2005). Fisher’s microscope and Haldane’s ellipse. The American Naturalist, 166: 447–457.

Wei, X. and Zhang, J. (2017). Why phenotype robustness promotes phenotype evolvability. Genome Biology and Evolution, 9: 3509–3515.

Wilensky, U. (1999). NetLogo. Center for Connected Learning and Computer-Based Modeling, Northwestern University, Evanston, IL, USA. http://ccl.northwestern.edu/netlogo.

Zheng, J., Payne, J. L. and Wagner, A. (2019). Cryptic genetic variation accelerates evolution by opening access to diverse adaptive peaks. Science, 365: 347–353.

