## Supplementary material for "A heuristic model of the effects of phenotypic robustness in adaptive evolution": Simulation scripts: Rigato and Fusco SIM 1-2.nlogo

```
turtles-own [
  fitness
  robustness
  ]
globals [ 
  carrying-capacity
  optimal-fitness
  count-ticks
  ]

to setup
  clear-all
  create-turtles population-size [
    set fitness 10
    set color fitness
    setxy random-xcor random-ycor
    set robustness phenotype-robustness 
    ]
  set carrying-capacity population-size
  set optimal-fitness 30
  set count-ticks 0 
  reset-ticks
end

to go
  if mean [fitness] of turtles > 30 [stop]
  ask turtles [
    move
    reproduce
    die
    ]
  ask n-of (0.1 * count turtles) turtles [die]
  
  
  tick
end

to reproduce
   if (count turtles < carrying-capacity) [
   hatch (fitness) [
      let genotype-mutation random-float 1 < gmp
      if genotype-mutation = true[
      let mutation random-float 1 < (1 - robustness)
      if mutation = true [
         mutate-fitness
        ]
      ]

      move
      ]
   ]
    
end

to move
  set heading random 360
    forward 2
end


to mutate-fitness
  let mutationsign random-float 1 < 0.17
  ifelse mutationsign = true [
    set fitness (mean [fitness] of turtles + s * mean [fitness] of turtles)] 
  [set fitness (mean [fitness] of turtles - s * mean [fitness] of turtles)]
   set color fitness 
end
@#$#@#$#@
GRAPHICS-WINDOW
739
10
1178
470
16
16
13.0
1
10
1
1
1
0
1
1
1
-16
16
-16
16
1
1
1
ticks
30.0

SLIDER
16
64
188
97
population-size
population-size
0
10000
1000
1
1
NIL
HORIZONTAL

SLIDER
16
106
232
139
phenotype-robustness
phenotype-robustness
0
1
0.5
0.0001
1
NIL
HORIZONTAL

PLOT
16
193
216
343
mean-fitness
time
NIL
0.0
10.0
0.0
1.0
true
false
"" ""
PENS
"default" 1.0 0 -16777216 true "" "plot mean [fitness] of turtles / optimal-fitness"

BUTTON
16
20
82
53
Setup
setup
NIL
1
T
OBSERVER
NIL
NIL
NIL
NIL
1

BUTTON
99
21
162
54
Go
go
T
1
T
OBSERVER
NIL
NIL
NIL
NIL
1

PLOT
228
195
435
346
mean-robustness
time
NIL
0.0
10.0
0.0
1.0
true
false
"" ""
PENS
"default" 1.0 0 -16777216 true "" "plot mean [robustness] of turtles"

MONITOR
449
300
535
345
Robustness
mean [robustness] of turtles
17
1
11

MONITOR
448
248
543
293
mean-fitness
mean [fitness] of turtles / optimal-fitness
17
1
11

MONITOR
447
196
558
241
population-size
count turtles
17
1
11

PLOT
16
350
216
500
variance-fitness
time
NIL
0.0
10.0
0.0
10.0
true
false
"" ""
PENS
"default" 1.0 0 -16777216 true "" "plot variance [fitness] of turtles"

MONITOR
450
354
629
399
fitness-variance
variance [fitness / 25] of turtles
17
1
11

MONITOR
449
409
657
454
robustness-variance
variance [robustness] of turtles
17
1
11

PLOT
227
355
427
505
robustness-variance
NIL
NIL
0.0
10.0
0.0
0.1
true
false
"" ""
PENS
"default" 1.0 0 -16777216 true "" "plot variance [robustness] of turtles"

SLIDER
290
95
462
128
gmp
gmp
0
1
0.5
0.01
1
NIL
HORIZONTAL

SLIDER
295
38
467
71
s
s
0
2
0.1
0.001
1
NIL
HORIZONTAL

@#$#@#$#@
## WHAT IS IT?

(a general understanding of what the model is trying to show or explain)

## HOW IT WORKS

(what rules the agents use to create the overall behavior of the model)

## HOW TO USE IT

(how to use the model, including a description of each of the items in the Interface tab)

## THINGS TO NOTICE

(suggested things for the user to notice while running the model)

## THINGS TO TRY

(suggested things for the user to try to do (move sliders, switches, etc.) with the model)

## EXTENDING THE MODEL

(suggested things to add or change in the Code tab to make the model more complicated, detailed, accurate, etc.)

## NETLOGO FEATURES

(interesting or unusual features of NetLogo that the model uses, particularly in the Code tab; or where workarounds were needed for missing features)

## RELATED MODELS

(models in the NetLogo Models Library and elsewhere which are of related interest)

## CREDITS AND REFERENCES

(a reference to the model's URL on the web if it has one, as well as any other necessary credits, citations, and links)
@#$#@#$#@
default
true
0
Polygon -7500403 true true 150 5 40 250 150 205 260 250

airplane
true
0
Polygon -7500403 true true 150 0 135 15 120 60 120 105 15 165 15 195 120 180 135 240 105 270 120 285 150 270 180 285 210 270 165 240 180 180 285 195 285 165 180 105 180 60 165 15

arrow
true
0
Polygon -7500403 true true 150 0 0 150 105 150 105 293 195 293 195 150 300 150

box
false
0
Polygon -7500403 true true 150 285 285 225 285 75 150 135
Polygon -7500403 true true 150 135 15 75 150 15 285 75
Polygon -7500403 true true 15 75 15 225 150 285 150 135
Line -16777216 false 150 285 150 135
Line -16777216 false 150 135 15 75
Line -16777216 false 150 135 285 75

bug
true
0
Circle -7500403 true true 96 182 108
Circle -7500403 true true 110 127 80
Circle -7500403 true true 110 75 80
Line -7500403 true 150 100 80 30
Line -7500403 true 150 100 220 30

butterfly
true
0
Polygon -7500403 true true 150 165 209 199 225 225 225 255 195 270 165 255 150 240
Polygon -7500403 true true 150 165 89 198 75 225 75 255 105 270 135 255 150 240
Polygon -7500403 true true 139 148 100 105 55 90 25 90 10 105 10 135 25 180 40 195 85 194 139 163
Polygon -7500403 true true 162 150 200 105 245 90 275 90 290 105 290 135 275 180 260 195 215 195 162 165
Polygon -16777216 true false 150 255 135 225 120 150 135 120 150 105 165 120 180 150 165 225
Circle -16777216 true false 135 90 30
Line -16777216 false 150 105 195 60
Line -16777216 false 150 105 105 60

car
false
0
Polygon -7500403 true true 300 180 279 164 261 144 240 135 226 132 213 106 203 84 185 63 159 50 135 50 75 60 0 150 0 165 0 225 300 225 300 180
Circle -16777216 true false 180 180 90
Circle -16777216 true false 30 180 90
Polygon -16777216 true false 162 80 132 78 134 135 209 135 194 105 189 96 180 89
Circle -7500403 true true 47 195 58
Circle -7500403 true true 195 195 58

circle
false
0
Circle -7500403 true true 0 0 300

circle 2
false
0
Circle -7500403 true true 0 0 300
Circle -16777216 true false 30 30 240

cow
false
0
Polygon -7500403 true true 200 193 197 249 179 249 177 196 166 187 140 189 93 191 78 179 72 211 49 209 48 181 37 149 25 120 25 89 45 72 103 84 179 75 198 76 252 64 272 81 293 103 285 121 255 121 242 118 224 167
Polygon -7500403 true true 73 210 86 251 62 249 48 208
Polygon -7500403 true true 25 114 16 195 9 204 23 213 25 200 39 123

cylinder
false
0
Circle -7500403 true true 0 0 300

dot
false
0
Circle -7500403 true true 90 90 120

face happy
false
0
Circle -7500403 true true 8 8 285
Circle -16777216 true false 60 75 60
Circle -16777216 true false 180 75 60
Polygon -16777216 true false 150 255 90 239 62 213 47 191 67 179 90 203 109 218 150 225 192 218 210 203 227 181 251 194 236 217 212 240

face neutral
false
0
Circle -7500403 true true 8 7 285
Circle -16777216 true false 60 75 60
Circle -16777216 true false 180 75 60
Rectangle -16777216 true false 60 195 240 225

face sad
false
0
Circle -7500403 true true 8 8 285
Circle -16777216 true false 60 75 60
Circle -16777216 true false 180 75 60
Polygon -16777216 true false 150 168 90 184 62 210 47 232 67 244 90 220 109 205 150 198 192 205 210 220 227 242 251 229 236 206 212 183

fish
false
0
Polygon -1 true false 44 131 21 87 15 86 0 120 15 150 0 180 13 214 20 212 45 166
Polygon -1 true false 135 195 119 235 95 218 76 210 46 204 60 165
Polygon -1 true false 75 45 83 77 71 103 86 114 166 78 135 60
Polygon -7500403 true true 30 136 151 77 226 81 280 119 292 146 292 160 287 170 270 195 195 210 151 212 30 166
Circle -16777216 true false 215 106 30

flag
false
0
Rectangle -7500403 true true 60 15 75 300
Polygon -7500403 true true 90 150 270 90 90 30
Line -7500403 true 75 135 90 135
Line -7500403 true 75 45 90 45

flower
false
0
Polygon -10899396 true false 135 120 165 165 180 210 180 240 150 300 165 300 195 240 195 195 165 135
Circle -7500403 true true 85 132 38
Circle -7500403 true true 130 147 38
Circle -7500403 true true 192 85 38
Circle -7500403 true true 85 40 38
Circle -7500403 true true 177 40 38
Circle -7500403 true true 177 132 38
Circle -7500403 true true 70 85 38
Circle -7500403 true true 130 25 38
Circle -7500403 true true 96 51 108
Circle -16777216 true false 113 68 74
Polygon -10899396 true false 189 233 219 188 249 173 279 188 234 218
Polygon -10899396 true false 180 255 150 210 105 210 75 240 135 240

house
false
0
Rectangle -7500403 true true 45 120 255 285
Rectangle -16777216 true false 120 210 180 285
Polygon -7500403 true true 15 120 150 15 285 120
Line -16777216 false 30 120 270 120

leaf
false
0
Polygon -7500403 true true 150 210 135 195 120 210 60 210 30 195 60 180 60 165 15 135 30 120 15 105 40 104 45 90 60 90 90 105 105 120 120 120 105 60 120 60 135 30 150 15 165 30 180 60 195 60 180 120 195 120 210 105 240 90 255 90 263 104 285 105 270 120 285 135 240 165 240 180 270 195 240 210 180 210 165 195
Polygon -7500403 true true 135 195 135 240 120 255 105 255 105 285 135 285 165 240 165 195

line
true
0
Line -7500403 true 150 0 150 300

line half
true
0
Line -7500403 true 150 0 150 150

pentagon
false
0
Polygon -7500403 true true 150 15 15 120 60 285 240 285 285 120

person
false
0
Circle -7500403 true true 110 5 80
Polygon -7500403 true true 105 90 120 195 90 285 105 300 135 300 150 225 165 300 195 300 210 285 180 195 195 90
Rectangle -7500403 true true 127 79 172 94
Polygon -7500403 true true 195 90 240 150 225 180 165 105
Polygon -7500403 true true 105 90 60 150 75 180 135 105

plant
false
0
Rectangle -7500403 true true 135 90 165 300
Polygon -7500403 true true 135 255 90 210 45 195 75 255 135 285
Polygon -7500403 true true 165 255 210 210 255 195 225 255 165 285
Polygon -7500403 true true 135 180 90 135 45 120 75 180 135 210
Polygon -7500403 true true 165 180 165 210 225 180 255 120 210 135
Polygon -7500403 true true 135 105 90 60 45 45 75 105 135 135
Polygon -7500403 true true 165 105 165 135 225 105 255 45 210 60
Polygon -7500403 true true 135 90 120 45 150 15 180 45 165 90

sheep
false
15
Circle -1 true true 203 65 88
Circle -1 true true 70 65 162
Circle -1 true true 150 105 120
Polygon -7500403 true false 218 120 240 165 255 165 278 120
Circle -7500403 true false 214 72 67
Rectangle -1 true true 164 223 179 298
Polygon -1 true true 45 285 30 285 30 240 15 195 45 210
Circle -1 true true 3 83 150
Rectangle -1 true true 65 221 80 296
Polygon -1 true true 195 285 210 285 210 240 240 210 195 210
Polygon -7500403 true false 276 85 285 105 302 99 294 83
Polygon -7500403 true false 219 85 210 105 193 99 201 83

square
false
0
Rectangle -7500403 true true 30 30 270 270

square 2
false
0
Rectangle -7500403 true true 30 30 270 270
Rectangle -16777216 true false 60 60 240 240

star
false
0
Polygon -7500403 true true 151 1 185 108 298 108 207 175 242 282 151 216 59 282 94 175 3 108 116 108

target
false
0
Circle -7500403 true true 0 0 300
Circle -16777216 true false 30 30 240
Circle -7500403 true true 60 60 180
Circle -16777216 true false 90 90 120
Circle -7500403 true true 120 120 60

tree
false
0
Circle -7500403 true true 118 3 94
Rectangle -6459832 true false 120 195 180 300
Circle -7500403 true true 65 21 108
Circle -7500403 true true 116 41 127
Circle -7500403 true true 45 90 120
Circle -7500403 true true 104 74 152

triangle
false
0
Polygon -7500403 true true 150 30 15 255 285 255

triangle 2
false
0
Polygon -7500403 true true 150 30 15 255 285 255
Polygon -16777216 true false 151 99 225 223 75 224

truck
false
0
Rectangle -7500403 true true 4 45 195 187
Polygon -7500403 true true 296 193 296 150 259 134 244 104 208 104 207 194
Rectangle -1 true false 195 60 195 105
Polygon -16777216 true false 238 112 252 141 219 141 218 112
Circle -16777216 true false 234 174 42
Rectangle -7500403 true true 181 185 214 194
Circle -16777216 true false 144 174 42
Circle -16777216 true false 24 174 42
Circle -7500403 false true 24 174 42
Circle -7500403 false true 144 174 42
Circle -7500403 false true 234 174 42

turtle
true
0
Polygon -10899396 true false 215 204 240 233 246 254 228 266 215 252 193 210
Polygon -10899396 true false 195 90 225 75 245 75 260 89 269 108 261 124 240 105 225 105 210 105
Polygon -10899396 true false 105 90 75 75 55 75 40 89 31 108 39 124 60 105 75 105 90 105
Polygon -10899396 true false 132 85 134 64 107 51 108 17 150 2 192 18 192 52 169 65 172 87
Polygon -10899396 true false 85 204 60 233 54 254 72 266 85 252 107 210
Polygon -7500403 true true 119 75 179 75 209 101 224 135 220 225 175 261 128 261 81 224 74 135 88 99

wheel
false
0
Circle -7500403 true true 3 3 294
Circle -16777216 true false 30 30 240
Line -7500403 true 150 285 150 15
Line -7500403 true 15 150 285 150
Circle -7500403 true true 120 120 60
Line -7500403 true 216 40 79 269
Line -7500403 true 40 84 269 221
Line -7500403 true 40 216 269 79
Line -7500403 true 84 40 221 269

wolf
false
0
Polygon -16777216 true false 253 133 245 131 245 133
Polygon -7500403 true true 2 194 13 197 30 191 38 193 38 205 20 226 20 257 27 265 38 266 40 260 31 253 31 230 60 206 68 198 75 209 66 228 65 243 82 261 84 268 100 267 103 261 77 239 79 231 100 207 98 196 119 201 143 202 160 195 166 210 172 213 173 238 167 251 160 248 154 265 169 264 178 247 186 240 198 260 200 271 217 271 219 262 207 258 195 230 192 198 210 184 227 164 242 144 259 145 284 151 277 141 293 140 299 134 297 127 273 119 270 105
Polygon -7500403 true true -1 195 14 180 36 166 40 153 53 140 82 131 134 133 159 126 188 115 227 108 236 102 238 98 268 86 269 92 281 87 269 103 269 113

x
false
0
Polygon -7500403 true true 270 75 225 30 30 225 75 270
Polygon -7500403 true true 30 75 75 30 270 225 225 270

@#$#@#$#@
NetLogo 5.0.3
@#$#@#$#@
@#$#@#$#@
@#$#@#$#@
<experiments>
  <experiment name="experiment" repetitions="3" runMetricsEveryStep="true">
    <setup>setup</setup>
    <go>go</go>
    <timeLimit steps="100"/>
    <metric>count turtles</metric>
    <metric>selection-differential</metric>
    <metric>count turtles</metric>
    <metric>mean [fitness / 30] of turtles</metric>
    <metric>mean [robustness] of turtles</metric>
    <metric>variance [fitness / 30] of turtles</metric>
    <metric>variance [robustness] of turtles</metric>
    <enumeratedValueSet variable="gmp">
      <value value="0.9"/>
    </enumeratedValueSet>
    <enumeratedValueSet variable="population-size">
      <value value="50"/>
      <value value="1000"/>
    </enumeratedValueSet>
    <enumeratedValueSet variable="phenotype-robustness">
      <value value="0.2"/>
      <value value="0.4"/>
      <value value="0.6"/>
      <value value="0.8"/>
      <value value="0.99"/>
    </enumeratedValueSet>
  </experiment>
  <experiment name="fixedsfixedrobustness" repetitions="3" runMetricsEveryStep="true">
    <setup>setup</setup>
    <go>go</go>
    <timeLimit steps="500"/>
    <metric>count turtles</metric>
    <metric>mean [fitness] of turtles / 30</metric>
    <enumeratedValueSet variable="population-size">
      <value value="1000"/>
      <value value="500"/>
      <value value="250"/>
      <value value="100"/>
      <value value="50"/>
      <value value="20"/>
    </enumeratedValueSet>
    <enumeratedValueSet variable="phenotype-robustness">
      <value value="0.7"/>
      <value value="0.75"/>
      <value value="0.8"/>
      <value value="0.85"/>
      <value value="0.9"/>
      <value value="0.95"/>
      <value value="0.99"/>
    </enumeratedValueSet>
    <enumeratedValueSet variable="s">
      <value value="0.1"/>
    </enumeratedValueSet>
    <enumeratedValueSet variable="gmp">
      <value value="1"/>
    </enumeratedValueSet>
  </experiment>
  <experiment name="experiment" repetitions="3" runMetricsEveryStep="true">
    <setup>setup</setup>
    <go>go</go>
    <timeLimit steps="100"/>
    <metric>count turtles</metric>
    <metric>mean [fitness] of turtles / optimal-fitness</metric>
    <enumeratedValueSet variable="population-size">
      <value value="1000"/>
    </enumeratedValueSet>
    <enumeratedValueSet variable="s">
      <value value="0.1"/>
    </enumeratedValueSet>
    <enumeratedValueSet variable="gmp">
      <value value="1"/>
    </enumeratedValueSet>
    <enumeratedValueSet variable="phenotype-robustness">
      <value value="0.3"/>
      <value value="0.4"/>
      <value value="0.5"/>
      <value value="0.6"/>
      <value value="0.7"/>
      <value value="0.8"/>
      <value value="0.9"/>
      <value value="1"/>
    </enumeratedValueSet>
  </experiment>
  <experiment name="experimentXX" repetitions="10" runMetricsEveryStep="false">
    <setup>setup</setup>
    <go>go</go>
    <timeLimit steps="500"/>
    <metric>count turtles</metric>
    <metric>mean[fitness] of turtles</metric>
    <enumeratedValueSet variable="population-size">
      <value value="1000"/>
    </enumeratedValueSet>
    <enumeratedValueSet variable="phenotype-robustness">
      <value value="0.0010"/>
      <value value="0.1"/>
      <value value="0.2"/>
      <value value="0.3"/>
      <value value="0.4"/>
      <value value="0.5"/>
      <value value="0.6"/>
      <value value="0.7"/>
      <value value="0.8"/>
      <value value="0.85"/>
      <value value="0.9"/>
      <value value="0.95"/>
      <value value="0.99"/>
    </enumeratedValueSet>
    <enumeratedValueSet variable="s">
      <value value="0.0010"/>
      <value value="0.0050"/>
      <value value="0.01"/>
      <value value="0.02"/>
      <value value="0.04"/>
      <value value="0.07"/>
      <value value="0.1"/>
    </enumeratedValueSet>
    <enumeratedValueSet variable="gmp">
      <value value="0.0010"/>
      <value value="0.1"/>
      <value value="0.2"/>
      <value value="0.3"/>
      <value value="0.4"/>
      <value value="0.5"/>
      <value value="0.6"/>
      <value value="0.7"/>
      <value value="0.8"/>
      <value value="0.9"/>
      <value value="0.99"/>
    </enumeratedValueSet>
  </experiment>
  <experiment name="experimentXXX" repetitions="10" runMetricsEveryStep="false">
    <setup>setup</setup>
    <go>go</go>
    <timeLimit steps="500"/>
    <metric>count turtles</metric>
    <metric>mean[fitness] of turtles</metric>
    <enumeratedValueSet variable="population-size">
      <value value="1000"/>
    </enumeratedValueSet>
    <enumeratedValueSet variable="phenotype-robustness">
      <value value="0.0010"/>
      <value value="0.1"/>
      <value value="0.2"/>
      <value value="0.3"/>
      <value value="0.4"/>
      <value value="0.5"/>
      <value value="0.6"/>
      <value value="0.7"/>
      <value value="0.8"/>
      <value value="0.85"/>
      <value value="0.9"/>
      <value value="0.95"/>
      <value value="0.99"/>
    </enumeratedValueSet>
    <enumeratedValueSet variable="s">
      <value value="0.1"/>
    </enumeratedValueSet>
    <enumeratedValueSet variable="gmp">
      <value value="0.0010"/>
      <value value="0.1"/>
      <value value="0.2"/>
      <value value="0.3"/>
      <value value="0.4"/>
      <value value="0.5"/>
      <value value="0.6"/>
      <value value="0.7"/>
      <value value="0.8"/>
      <value value="0.9"/>
      <value value="0.99"/>
    </enumeratedValueSet>
  </experiment>
</experiments>
@#$#@#$#@
@#$#@#$#@
default
0.0
-0.2 0 1.0 0.0
0.0 1 1.0 0.0
0.2 0 1.0 0.0
link direction
true
0
Line -7500403 true 150 150 90 180
Line -7500403 true 150 150 210 180

@#$#@#$#@
0
@#$#@#$#@
```
